# Reproductive Isolation through Experimental Manipulation of Sexually Antagonistic Coevolution in *Drosophila melanogaster*

**DOI:** 10.1101/079657

**Authors:** Syed Zeeshan Ali, Martik Chatterjee, Manas Arun Samant, Nagaraj Guru Prasad

## Abstract

Promiscuity can drive the evolution of sexual conflict before and after mating occurs. Post-mating, the male ejaculate can selfishly manipulate female physiology leading to a chemical arms race between the sexes. Theory suggests that drift and sexually antagonistic coevolution can cause allopatric populations to evolve different chemical interactions between the sexes, thereby leading to postmating reproductive barriers and speciation. There is, however, little empirical evidence supporting this form of speciation. We tested this theory by creating an experimental evolutionary model of *Drosophila melanogaster* populations undergoing different levels of interlocus sexual conflict. We found that allopatric populations under elevated sexual conflict show assortative mating indicating premating reproductive isolation. Further, these allopatric populations also show reduced copulation duration and sperm defense ability when mating happens between individuals between individuals across populations compared to that within the same population, indicating postmating prezygotic isolation. Sexual conflict can cause reproductive isolation in allopatric populations through the coevolution of chemical (postmating prezygotic) as well as behavioural (premating) interaction between the sexes. Thus, to our knowledge, we provide the first comprehensive evidence of postmating (as well as premating) reproductive isolation due to sexual conflict.

## Introduction

In sexually reproducing species, males and females often have differential reproductive investment, and, consequently differential evolutionary interest in the outcome of sexual interactions^1, 2^. This often leads to a scenario where adaptations benefitting one sex come at the expense of the other^3, 4, 5^, ensuing a coevolutionary chase typically called sexually antagonistic coevolution (SAC)^6^. According to verbal^7, 8^ and formal^9, 10^ arguments, SAC can lead to a perpetual arms race between males and females of the same species. A byproduct of this is the continual divergence between allopatric populations in genes related to reproduction, leading to reproductive isolation (RI) even in the absence of natural selection. This hypothesis is supported indirectly by comparative studies that showed higher rates of speciation in insect clades where sexual conflict is observed than those where it is not observed^11^. However, no such evidence is found in other studies on mammals, butterflies, spiders^12^ and birds^13^.

An alternative to phylogenetic analysis that has been used to directly test the hypothesis is through experimental evolution which generally follows a simple experimental design:

a. Evolving independent replicate (i.e., allopatric) populations maintained under high and low/no conflict regimes (e.g., by enforcing monogamy or altering sex ratio) while all else remains equal.
b. Thereafter quantifying RI between allopatric populations within a regime and comparing the extent of isolation between different regimes.

Following this hypothesis, upon secondary contact, allopatric populations will show relatively stronger evidence of RI in the high conflict regime compared to low/no conflict regime. Martin and Hosken tested the hypothesis in *Sepsis cynipsea* by evolving replicate populations under polygamy (SAC) and monogamy (removal of SAC) for 35 generations. They found that allopatric pairs showed significantly less mating success compared to their sympatric counterparts in the polygamous, but not in monogamous regime, thus providing the first evidence that antagonistically evolving behavioural traits can lead to reproductive isolation^14^.

Along with premating behavioural interactions, postmating chemical interactions are important players in driving SAC. Ejaculate-female interaction and subsequent coevolution has been shown to have caused diversification in both ejaculate components (e.g. sperm, accessory gland proteins, small molecules transferred through ejaculate) and female reproductive tract and behaviour across taxa^15^. Thus, postmating antagonistic coevolution can lead to postmating RI through an ‘assortative sperm/ejaculate choice’ process that is analogous to assortative mate choice. However, there is no empirical evidence favouring this. Despite multiple studies testing the hypothesis in different organisms, the study by Martin and Hosken remains the only direct evidence of SAC as a driver of RI so far^16, 17, 18, 19, 20^, and the idea of sexual conflict as an ‘engine of speciation’ remains controversial^21^.

We used two sets of allopatric populations of *Drosophila melanogaster* – one set (of three populations) evolving under male biased (M) operational sex ratio and the other set (of three populations) evolving under female biased (F) operational sex ratio, demonstrating high and low levels of SAC respectively^4, 5^. We tested whether reproductive isolation between allopatric populations was more prominent, if not present only in M as compared to F regime.

Reproductive isolation can manifest in three stages: premating, postmating prezygotic and postzygotic^22^. We have focused on the first two as they are expected to evolve rapidly and have higher probabilities of being manifested^7^ within the relatively shorter time scale of experimental evolution.

As a measure of premating isolation, we assayed (a) assortative mating between females and males from the same population in presence of a competitor male from a different population (within the same regime) and (b) female reluctance to mate. As for postmating prezygotic isolation, we compared (a) copulation duration and (b) competitive fertilisation success of males from within and across population crosses.

## Results

The selection lines were derived from a long term laboratory adapted population of *Drosophila melanogaster* called LH_st_^23^. The LH_st_ population was in turn derived by the introgression of an autosomal recessive ‘scarlet eye’ (st) mutation to another large laboratory bred population called LH (see methods for further description of ancestral populations).

Each of the three independent replicates of female biased regime (F_1,2,3_) and male biased regime(M_1,2,3_) were created by altering the sex ratio to female biased (1 male:3 female) and male biased (3 male: 1 female) respectively^24^. All assays were done between the 95^th^ and 105^th^ generations of selection.

Males and females used in the assays were either from the same replicate population or from different replicate populations within a regime, which we term as ‘within replicate’ (WR) and ‘between replicate’ (BR) respectively. Flies used for all the assays were collected as virgins and a held singly in vials (90-mm length × 30-mm diameter) containing fresh corn meal - yeast-molasses food. All flies were 2-3 day-old adults at the time of assay.

### Assay for premating isolation

To look for premating reproductive isolation though assortative mating, we combined a virgin female with a WR and a BR virgin male (simultaneously) in a round-robin manner and observed which one of the two males mated with the female (Table 1). A logistic regression showed higher degree of assortative mating in M regime compared to that in F, i.e., in M regime, WR males mated more successfully compared to BR males (p= 0.0488). While the log-odds ratio showed no difference in choice for F regime (“intercept” in Table 2), in M regime the ratio increases significantly for successful WR matings in M regime (“SelectionM” in table 2). This suggests premating reproductive isolation between allopatric populations in the M regime. However, another measure of premating isolation, mating latency (time taken for a pair to start mating after they are combined) showed no evidence of reproductive isolation (two way ANOVA: F_1,221_=1.678, p = 0.613, Figure 1).

**Figure 1:**
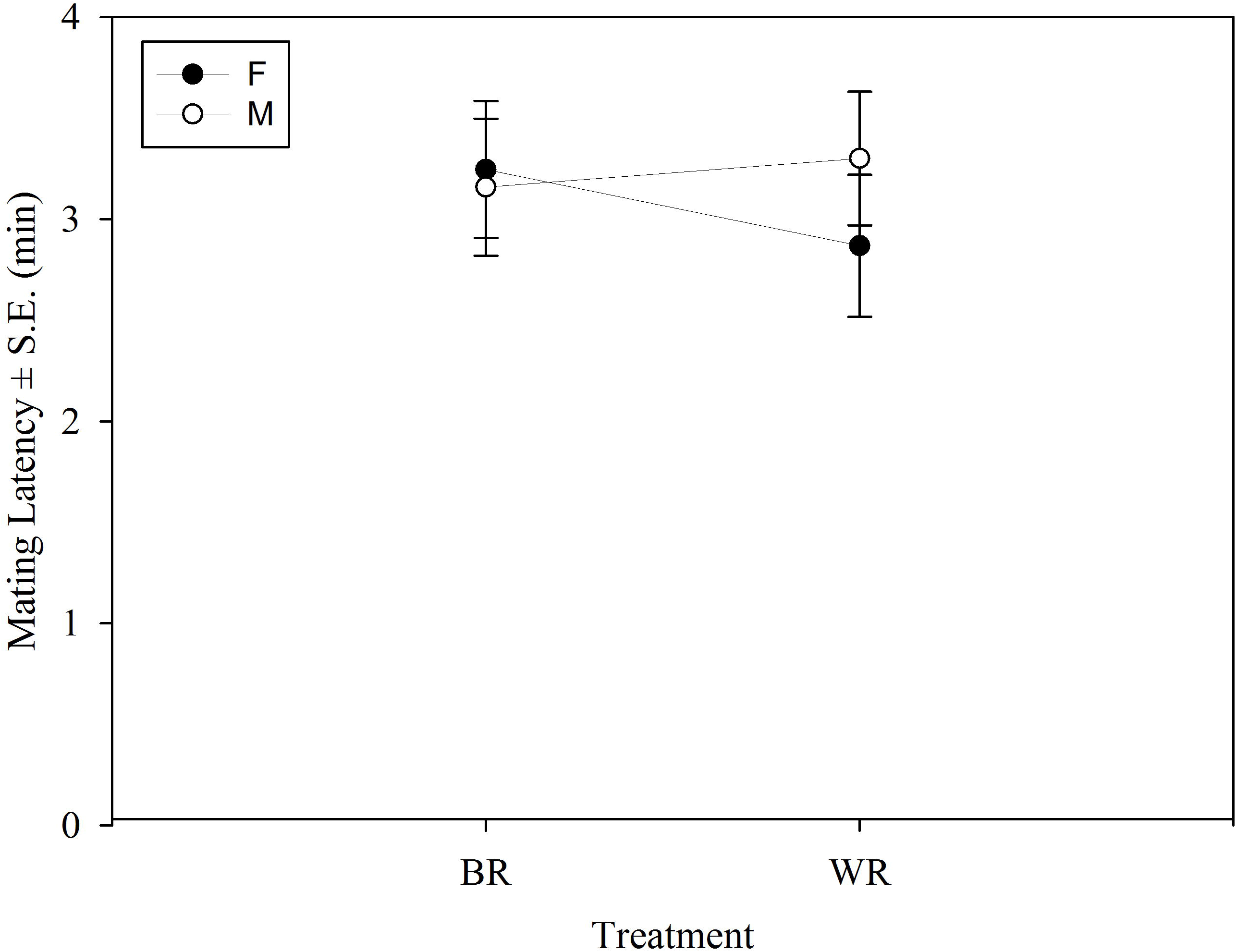
Mean mating latency (±S.E) of WR and BR treatments from female biased (F) and male biased (M) regimes based on the results of two-way ANOVA. There was no significant Selection Regime × Treatment interaction.

**Table 1:** Mating treatments for different assays. The letters i and j denote block (replicate) numbers, i **≠** j (in a round robin way). All mating trials were conducted within a selection regime.

| Assay | Female from block | Male from block | Sample size |
| --- | --- | --- | --- |
| Assortative mating | i | i(pink) + j(green) | 30 |
|  |  | j(green) +i(pink) | 30 |
| Mating latency, copulation duration | i | i | 20 |
| Sperm defence ability | i | j | 30 |

**Table 2.** Results of logistic regressionperformed on the data obtained from the assay for assortative mating. Successful mating between WR individuals was used as the response variable. Selection regime was used as fixed effect and replicate population nested within Selection regime was used as random effect.

|  | Estimate | Std. Error | z value | Pr(>\|z\|) |
| --- | --- | --- | --- | --- |
| Intercept | -0.0470675 | 0.15343501 | -0.3067586 | 0.759027 |
| SelectionM | 0.43273 | 0.21965409 | 1.9700521 | 0.048832 |

### Assay for postmating prezygotic isolation

To test for postmating prezygotic isolation, we first measured copulation duration (the time spent *in-copula* by a mating pair). Within each selection regime we had two treatments where one virgin female was combined with either one virgin BR or one virgin WR male. We had 60 replicate vials per treatment (WR/BR) per selection regime (M/F) for this experiment (Table 1).

In a two way ANOVA using treatment and selection as fixed factors, we found a significant selection regime × treatment interaction (F_1, 221_= 4.269, p = 0.03, Figure 2). Tukey’s HSD showed that in F there was no difference in copulation duration but in M, copulation duration was significantly higher in WR crosses compared to BR crosses (Table 3).

**Figure 2:**
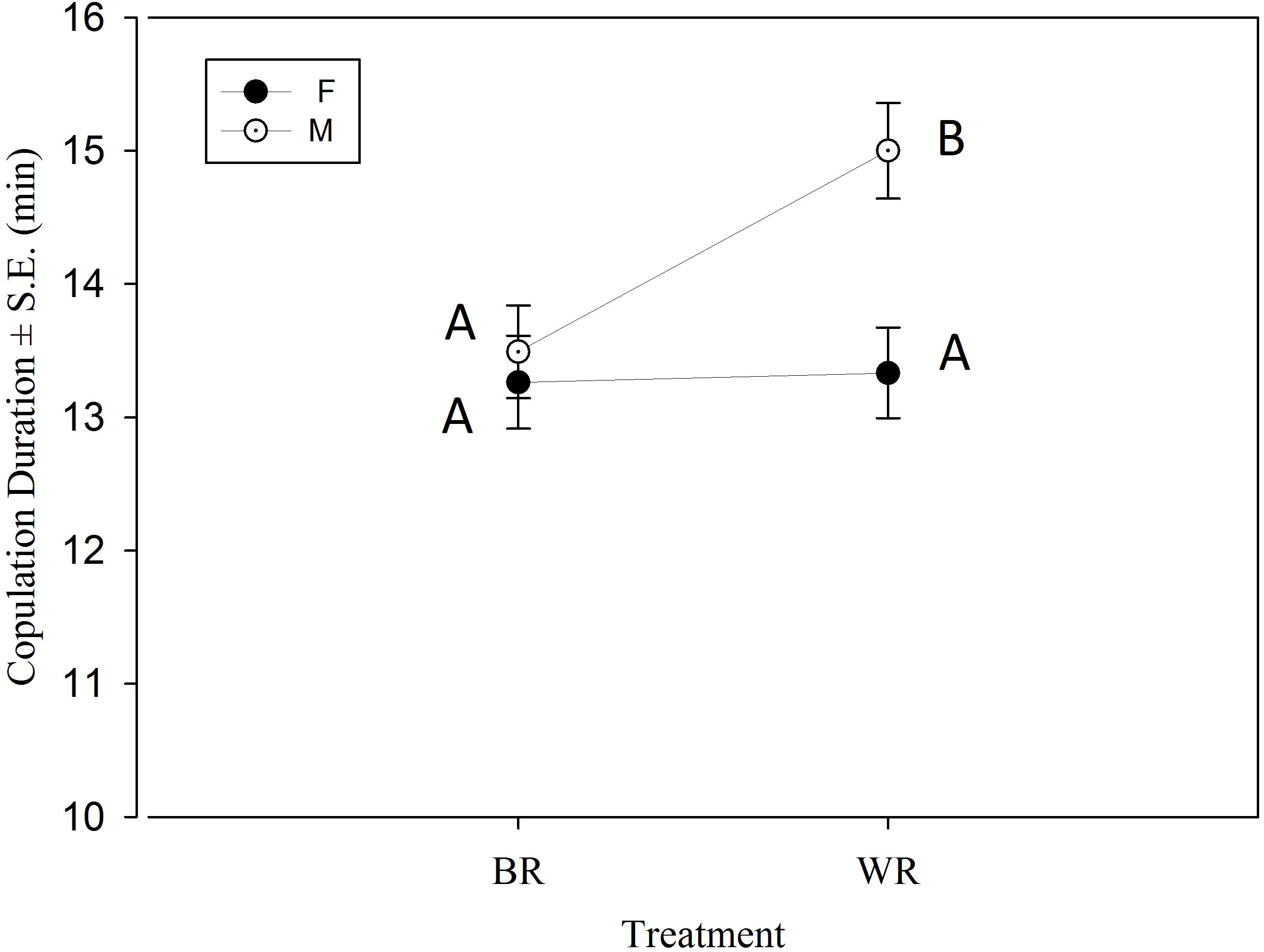
Mean copulation duration (±S.E) of WR and BR treatments from female biased (F) and male biased (M) regimes based on the results of two-way ANOVA. Points not sharing common letter (e.g., A and B) are significantly different based on Tukey’s HSD.

**Table 3.** A. Least square means ± S.E. values for Copulation Duration and P1. B. P values obtained through pairwise contrast using Tukey’s HSD. The important contrasts have been highlighted in bold.

**B. P values obtained through pairwise contrast using Tukey's HSD. The important contrasts have been highlighted in bold.**
| A. Lsmeans |  |  |  |
| --- | --- | --- | --- |
| Selection Regime | Mating Type | Lsmeans ± S.E. | |
|  |  | Copulation Duration | ArcSineSqrt (P1) |
| F | BR | 13.2359 ± 0.688966 | 0.349871 ± 0.0504 |
| M | BR | 13.47331 ± 0.688967 | 0.329624 ± 0.051427 |
| F | WR | 13.33333 ± 0.68479 | 0.373822 ± 0.050413 |
| M | WR | 15.01464 ± 0.695161 | 0.498448 ± 0.053216 |
| B. Contrasts (Tukey's HSD) |  |  |  |
| Contrast | p.values |  |  |
|  | Copulation Duration | AecSineSqrt (P1) |  |
| F BR - M BR | 0.959157 | 0.97628 |  |
| **F BR - F WR** | **0.996832** | **0.959197** |  |
| F BR - M WR | 0.001716 | 0.019729 |  |
| M BR - F WR | 0.990766 | 0.804703 |  |
| **M BR - M WR** | **0.009053** | **0.007008** |  |
| F WR - M WR | 0.003004 | 0.070632 |  |

The difference in copulation duration was an indication of incipient reproductive isolation in terms of reproductive behaviour. We have previous evidence that in the ancestral population, copulation duration of the first mating is positively correlated with sperm defense ability^25^. So we tested if such a behavioural change translates into fitness difference. Sperm defense ability (P1) is measured as the proportion of progeny sired by the first male when the female is mated with multiple males (typically two males for assay purposes). A two way ANOVA similar to that of mating latency and copulation duration showed a significant selection regime × treatment interaction (F_1, 306_=4.198, p = 0.041, Figure 3). Tukey’s HSD and showed that in F, P1 of WR and BR males were not different but in M, WR males had significantly higher P1 value compared to that of the males from BR crosses (Table 3). This indicates that the difference in mating behaviour also translates into fitness differences.

**Figure 3:**
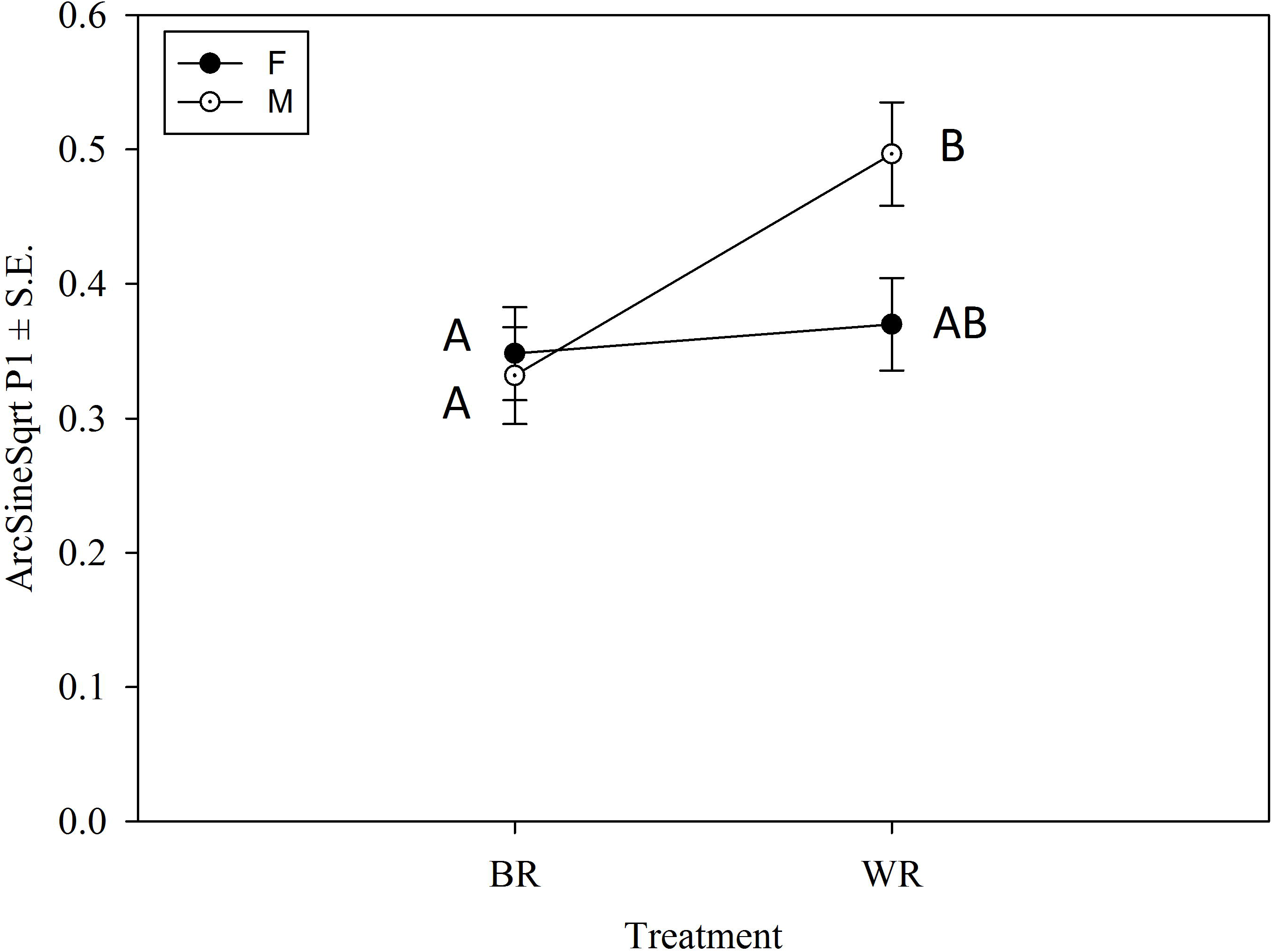
Mean (arcsine square root transformed) P1 (±S.E) of WR and BR treatments from female biased (F) and male biased (M) regimes based on the results of two-way ANOVA. Points not sharing common letter (e.g., A and B) are significantly different based on Tukey’s HSD.

## Discussion

In this study, we used experimental evolution to show that high levels of SAC can lead to the evolution of early stages of reproductive isolation at (a) premating and (b) postmating prezygotic stages in populations of *Drosophila melanogaster*.

In populations under high sexual conflict (M), females mated primarily with males of the same population in presence of an allopatric competitor from the same regime. Populations under low sexual conflict (F), on the other hand, displayed no such trend. Our observations corroborate that of Martin and Hosken^14^, who found evidences of premating isolation in dung fly populations maintained under promiscuous (but not monogamous) conditions. However, unlike them, we did not find any difference in females’ reluctance to mating (measured as mating latency in our study) under non-competitive scenario. The most plausible reason for this is the interspecies difference of the two studies. In Sepsis flies used in the previous study^14^, females show conspicuous reluctance by shaking and kicking. While such conspicuous behaviour is not present in *Drosophila melanogaster*, we found that premating isolation can manifest under a choice scenario. In the M populations SAC might have created genetic divergence, due to which females get spotted faster and/or courted more vigorously by WR males or simply find WR males more attractive than their BR counterparts leading to assortative mating. Thus, we provide evidence that premating RI can manifest itself under competitive scenario in terms of *mate choice* behaviour in addition to/instead of failed mating or ‘reluctance to mate’ behaviour – a possibility that has largely been neglected by most previous studies^16, 17, 18, 19^. However, Plesnar-Bielak *et al*. address this possibility but find no effect of SAC on assortative mating in the bulb mite *Rhizoglyphus robini*, after maintaining them under monogamous or polygamous regimes for 45 generations^20^.

Our assays resulted in WR pairs mating for longer and males enjoying greater sperm defense ability (when competed with common baseline males) than their BR counterparts in M populations but not in F. Thus in these populations, SAC seems to have resulted in postmating prezygotic RI between allopatric populations.

Copulation duration is an important indicator of male ejaculate investment as well as cryptic male mate choice^30, 31^. In a similar study on *Drosophila pseudoobscura*, Bacigalupe *et al.* used copulation duration as a one of the measures of reproductive isolation. In that, they evolved populations under different intensities of SAC and compared difference in copulation duration (among other traits) between WR and BR crosses. They found significant difference only in the regime with the highest SAC intensity, where WR crosses had lower copulation duration than BR crosses^18^. Our result is in stark contradiction to that. Copulation duration has also been used as an indicator of reproductive isolation in speciation studies on several *Drosophila* species complexes^26, 27, 28^. In all the studies, individuals from sister species did mate, but at least in some cases, heterospecific matings had lower copulation duration than conspecific matings. Our results could represent an early stage of speciation in this regard. Lower copulation duration in BR mating compared to WR mating in M populations could be due to genetic divergence caused by SAC that leads to reduced ejaculate transfer ability and/or cryptic male investment by the males when they mate with allopatric females.

A number of studies - while testing if SAC drives reproductive isolation using experimental evolution - have measured postmating isolation extensively in terms of difference in fecundity^17^, offspring number^19, 20^, offspring viability^17, 18^ or offspring sterility^18^, but found no evidence of isolation in those traits. While important, most of them are measures of post-zygotic isolation, which, as these and other studies^7^ suggest, are less likely to manifest within tens of generations of selection. Therefore, we focused primarily on prezygotic isolation. An important measure of prezygotic isolation is competitive fertilisation success^8^ which none of the studies thus far has addressed. We found that M males have lower competitive fertilisation success when competition happens in BR females than when it does in WR females, while in F males there is no such difference. Since in these populations we could not assay sperm competition directly between BR and WR males due to the lack of phenotypic markers, we have used a proxy measure where all the competitor males used in these assays were taken from the same ancestor population with the assumption that relative sperm competitive ability of the common competitors do not differ across replicate populations within a regime. This is a valid assumption since in a previous study comparing sperm competitive ability of M and F males (where we used the same common competitors) we found no replicate effect ^23^.

There are at least two reasons why M males have reduced sperm competitive ability when mated with allopatric M females. First, it could be a direct correlate of decreased copulation duration. Males with lower copulation duration do not/cannot transfer as much ejaculate and therefore have lower competitive ability^29^. The copulation duration-competitive ability correlation has been demonstrated in the ancestral population from which the selected populations have been derived^25^. Second, it could be a putative stage of conspecific sperm precedence (CSP) –where sperm of conspecific male has greater competitive success over that of heterospecific male. Evidence of CSP is widespread across various taxa^30, 31, 32, 33^ and its mechanisms have been illustrated for at least one set of Drosophila sibling species^32, 33^. In *Drosophila melanogaster* (as in most promiscuous species) females mate multiple times and often store ejaculate (in specialized storage organs, e.g., seminal receptacle and spermatheca in fruit flies) from different males where they compete for fertilisation success. The outcome is mostly determined by how the resident ejaculate (from an earlier mating) is displaced from female storage organs by ejaculate from more recent mating^34^ and is influenced by competing males and host female^35^. This provides ample scope for sperm-female coevolution^36^. Since at least some accessory gland proteins are harmful to females, ejaculate-female coevolution should be antagonistic in nature. It is possible that such postmating SAC drove divergence in replicate M populations in terms of how ejaculate and female reproductive tract interact to determine fertilisation success, leading to an incipient form of CSP. Thus, our results show higher rates of SAC can drive reproductive isolation in allopatric populations through reduced postmating competitive success of males.

Out of all the studies that have used experimental evolution to test the theoretical prediction that sexually antagonistic coevolution can drive reproductive isolation, there are only two (including the present one) that provide evidence in support, and to the best of our knowledge, this is the only one that provides evidence of postmating isolation. There are multiple reasons as to why our results differ from most of its predecessors^16, 17, 18, 19, 20^:

a. The census population size for each replicate was bigger in our study than those of the previous ones.
b. The number of generations in those studies were too low (our assays were done after ∼100 generations of selection compared to that of ≤ 50 in all of the previous studies) to allow SAC to drive population divergence to a degree where they are apparent.
c. According to theoretical predictions, reproductive isolation in allopatric populations is one of the six possible outcomes of sexual conflict^9^. It is possible that the populations under high SAC in those studies did not diverge with respect to each other. However, none of the studies shed light upon any of the other five possibilities that might have occurred in their populations.

In conclusion, we show direct evidence of evolution of both premating and postmating prezygotic RI as a consequence of SAC. Thus, it remains a distinct possibility that sexual conflict can result in a coevolutionary chase between the sexes^11, 37^ and can indeed be ‘an engine of speciation’. We speculate that initial genetic variation and number of generations can be important to realize – at least in experimental evolution studies –the evolution of RI caused by sexual conflict. However we also feel the need of more such studies to experimentally determine the exact conditions under which sexual conflict leads to reproductive isolation and to elucidate the underlying proximate mechanisms.

## Methods

### Ancestral Populations

LH – It is a large laboratory adapted population of *Drosophila melanogaster,* established by, and named after Lawrence G. Harshman. The population is maintained on a 14 day discrete generation cycle, under 25°C, 60-80% relative humidity, 12 hours light / 12 hours dark (12hrs: 12hrs L/D cycle) and on standard cornmeal – molasses – yeast food. The flies are grown under moderate larval density of 140-160 per 8-dram vial (25mm diameter × 90mm height) containing 8-10ml food. On the 12^th^ day post egg collection, flies from different vials are mixed and redistributed across fresh food vials containing limiting amount of live yeast grains with 16 males and 16 females per vial. On the 14^th^ day, flies are transferred to fresh vials and are allowed a window of 18 hours to lay eggs which (after discarding the adults and controlling density) start the next generation^38^.

LH_st_ – This population was derived by introducing the scarlet eye colour (recessive, autosomal and benign) gene into the LH population, hence the subscript. LH_st_ is maintained under the same condition as LH with N_e_>2500. The genetic backgrounds of these two populations are homogenized by periodic back crossing.

### Selection Regimes

The study was done on six populations of *Drosophila melanogaster* – M_1-3_ and F_1-3_ representing male biased and female biased operational sex ratio respectively. All these populations were created from the LH_st_ population.

We derived the male biased (M_1-3_) and female biased (F_1-3_) regimes, each having three independent replicates, from LH_st_ by varying the operational sex ratio to male: female:: 3:1 and 1:3 respectively. The maintenance of these populations differs from that of LH and LH_st_in the following ways:

a. In these populations adult flies are collected as virgins 9-10 days after egg collection, during the peak eclosion period and held in vials (containing 8 flies of one sex) for two days.
b. The sexes are combined on the 12^th^ day in fresh food vials seeded with measured amount of live yeast (0.47mg per female) following the selection regime – 24 males + 8 females in each vial for M and 8 males + 24 females in each vial for F.

The effective population sizes of all the populations are maintained at > 450 or > 350 depending on the method used to calculate them^4^. For more details on the evolutionary history and detailed maintenance protocol, see 23.

### Standardisation and Generation of Experimental Flies

In order to equalise the potential non-genetic parental effects across different regimes, we maintained all populations under ancestral condition which does not include virgin collection and sex ratio alteration-essentially following the same life cycle as LH_st_ populations for one generation before obtaining individuals for the experiment. This process is called standardisation^39^.

Eggs laid by the standardised flies were collected at a density of 150 (±2) per vial (containing 8-10ml of cornmeal food) to obtain the experimental flies. On the 10^th^ day after egg collection, males and females were collected as virgins during the peak of their eclosion and held as single individual per vial.

Ancestral flies (LH), whenever they were used in this study, were raised in similar conditions. LH males were sorted on the 12^th^ day post eclosion and held as single individuals. Eggs for LH flies were collected on the same day as that of the selection lines. Thus, the age of the experimental flies of all the populations was the same during the experiment.

### General Experimental Design

For all our assays, we compared reproductive behaviour and/or fitness related traits between two types of individuals within a regime:

a. Within replicate (WR): These are individuals from the same replicate number of a given selection regime i.e., M_i_♂ and M_i_♀ are WR with respect to each other where i denotes the replicate number (e.g., M_1_ ♂and ♀) and similarly for F.
b. Between replicate (BR): These are individuals from different replicate numbers of a given selection regime, i.e., M_i_♂ and M_j_♀ are BR with respect to each other –where i, j denote replicate numbers and (i,j) ͼ {(1,2), (2,3), (3,1)} (e.g. M_1_♂ and M_2_♀) and similarly for F. We took BR individuals in a round robin manner to avoid the problem of pseudo-replication^21^.

### Assay for Assortative Mating

We combined a virgin female with two virgin males from the same selection regime –one WR and one BR – in vials containing fresh food. That is, a female from a given replicate number was combined with a male from the same replicate number and another from a different replicate number (all within the same selection regime), e.g., one M_1_ female + one M_1_ male + one M_2_ male and so on. Thus, we had three combinations within each selection regime, denoted by female replicate number. Males were marked by pink or green Day-Glo dust for identification. Previous studies using the same dust found no effect on individuals in terms of mating behaviour or female preference ^40^. However, to account for any mating bias brought about solely by green and/or pink coloration, we had reverse coloration treatments for all combinations. Thus, each combination had two treatments, e.g., one M_1_ female + one green M_1_ male + one pink M_2_ male; one M_1_ female + one pink M_1_ male + one green M_2_ male and so on. We had 30 replicate vials per combination per colour treatment (Table1). In some vials we observed no mating till one hour after combining the flies. Those vials were discarded and excluded from analysis (Number of discarded vials: 4, 4, and 2 out of 60 trials each from the three replicates in the F regime; 4, 7, and 6 out of 60 trials each from the three replicates in the M regime).

### Assay for Mating Latency and Copulation Duration

For this assay we combined one virgin male and one virgin female according to treatment (WR or BR, see results) in a vial containing fresh food. After combining a male and a female, the pair was observed till they finished mating. Time taken for a pair to start mating after they were combined was recorded as mating latency and the time they spent in-copula was recorded as copulation duration. If a pair failed to mate after one hour, they were discarded. However, the number of failed mating in all treatments was very low (6, 3, 0 and 3 failures out of 60 trials in M-WR, M-BR, F-WR and F-BR respectively). Mating latency and copulation duration values for each vial were used as the unit of replication.

### Assay for Competitive fertilisation success

As a measure of competitive fertilisation success, we measured sperm defense ability of males, the rationale for which is provided in the results section. For assaying sperm defense ability, we set up crosses following the same method as mentioned above and the vials were observed for mating for one hour. The females that did not mate with the first male were discarded. After the first mating, we sorted the females using light CO_2_-anaesthesia and held them back into the vials and discarded the males. After allowing a recovery time (from anesthesia) of half an hour, we introduced a second male (red eyed, LH) in each vial and kept the vials undisturbed for 24 hours, during which they could mate with the females. After this exposure window, the second males were discarded and the females were transferred singly (under light anesthesia) to test tubes (dimensions: 12 mm diameter × 75 mm length) provisioned with food. There they were allowed an oviposition window of 18 hours. The adult progeny emerging from the eggs laid during this window were scored for their eye colour marker after 12 days. The proportion of scarlet progeny was taken as an estimate of P1 of the male. 90 males from each of the crosses were assayed for P1. Since we did not observe the second mating, instances where all progeny was sired only by the first male (P1 = 1) could arise due to second male failing to mate. Such instances were excluded from the analysis. Final sample size for P1 analysis was n = 83-87 and 70-73 per cross type (WR/BR) in F and M populations respectively. P1 value from a single vial was used as the unit of replication.

### Statistical Analysis

To test for assortative mating, we used logistic regression using a mixed model in a nested structure. In the model, the successful mating by a WR male was used as the response variable, selection regime was used as a fixed effect, and replicate population (to which the female belonged) nested within selection regime was used as a random effect using the following model:

WR_Success ∼ Selection +(Selection|Block), family = binomial(logtit)

The ‘glmer’ function in the ‘lme4’ package^42^ in R^43^ with binomial (logit) family was used for the analysis.

For the rest of the assays, we performed a two-way ANOVA with selection regime and treatment (type of individuals involved in a cross: BR/WR) as fixed factors and male and female replicate nested within selection regime as random effects using the following model:

response ∼ Sel.Reg*Mating.Type +(1|Female.Replicate) + (1|Male.Replicate)

In case of significant Sel.Reg*Mating.Type interaction, we performed Tukey’s HSD (with α = 0.05) for post-hoc analysis. Linear model fitting, ANOVA and post-hoc tests were performed in R using packages ‘lme4’, ‘lmerTest’^44^ and ‘lsmeans’^45^ respectively.

## Acknowledgement

The authors would like to thank Dr Bodhisatta Nandy, Dr Vanika Gupta, Ms Sharmi Sen, Prof. Amitabh Joshi, Prof. Adam K Chippindale, Mr Sudipta Tung, Dr Sutirth Dey and Dr TNC Vidya for helpful discussions and comments on the manuscript; Mr Tejinder Singh Chechi for help in population maintenance; numerous undergraduate and masters’ students of IISER Mohali for help in data collection.

## Author Contributions

SZA designed the study, carried out experiments, analysed data and wrote the manuscript. MC and MAS carried out experiments. NGP designed the study, and wrote the manuscript. All authors reviewed the manuscript.

## Additional information

### Competing financial interest

The authors declare that they have NO competing financial interest.

